# Identification of microRNAs that stabilize p53 in HPV-positive cancer cells

**DOI:** 10.1101/2020.09.21.305946

**Authors:** Gustavo Martínez-Noël, Patricia Szajner, Rebecca E. Kramer, Kathleen A. Boyland, Asma Sheikh, Jennifer A. Smith, Peter M. Howley

**Affiliations:** Department of Immunology, Harvard Medical School, 77 Avenue Louis Pasteur, Boston MA 02115; ICCB-Longwood Screening Facility, Harvard Medical School, 250 Longwood Avenue, Boston MA 02115

## Abstract

Etiologically, 5% of all cancers worldwide are caused by the high-risk human papillomaviruses (hrHPVs). These viruses encode two oncoproteins (E6 and E7) whose expression is required for cancer initiation and maintenance. Among their cellular targets are the p53 and the retinoblastoma tumor suppressor proteins. Inhibition of the hrHPV E6-mediated ubiquitylation of p53 through the E6AP ubiquitin ligase results in the stabilization of p53, leading to cellular apoptosis. We utilized a live cell high throughput screen to determine whether exogenous microRNA (miRNA) transfection had the ability to stabilize p53 in hrHPV-positive cervical cancer cells expressing a p53-fluorescent protein as an *in vivo* reporter of p53 stability. Among the miRNAs whose transfection resulted in the greatest p53 stabilization was 375-3p that has previously been reported to stabilize p53 in HeLa cells, providing validation of the screen. The top 32 miRNAs in addition to 375-3p were further assessed using a second cell-based p53 stability reporter system as well as in non-reporter HeLa cells to examine their effects on endogenous p53 protein levels, resulting in the identification of 23 miRNAs whose transfection increased p53 levels in HeLa cells. While a few miRNAs that stabilized p53 led to decreases in E6AP protein levels, all targeted HPV oncoprotein expression. We further examined subsets of these miRNAs for their abilities to induce apoptosis and determined whether it was p53-mediated. The introduction of specific miRNAs revealed surprisingly heterogeneous responses in different cell lines. Nonetheless, some of the miRNAs described here have potential as therapeutics for treating HPV-positive cancers.

**Importance:** Human papillomaviruses cause approximately 5% of all cancers worldwide and encode genes that contribute to both the initiation and maintenance of these cancers. The viral oncoprotein E6 is expressed in all HPV-positive cancers and functions by targeting the degradation of p53 through the engagement of the cellular ubiquitin ligase E6AP. Inhibiting the degradation of p53 leads to apoptosis in HPV-positive cancer cells. Using a high throughput live cell assay we identified several miRNAs whose transfection stabilize p53 in HPV-positive cells. These miRNAs have the potential to be used in the treatment of HPV-positive cancers.

## Introduction

Cervical cancer is the second leading cause of cancer deaths among women worldwide, with approximately 500,000 new cases diagnosed and 275,000 deaths each year. Virtually all cases of cervical cancer are attributable to infection by HPVs. There are over 200 different HPV types and a subset of these are linked to anogenital tract infections, 14 of which are referred to as hrHPV because they are associated with lesions that can progress to cancer (1). These same hrHPVs are also associated with other genital tract cancers (penile, vaginal, vulvar, and anal), as well as an increasing number of oropharyngeal cancers. Indeed, approximately 5% of all cancers worldwide are caused by hrHPVs (2).

Despite the success of the virus like particle (VLP)-based vaccine that has the potential to prevent millions of new HPV-associated cancers, there is the need for effective therapies to treat cancers that will develop in individuals who have not been vaccinated and those already infected with hrHPVs. In this regard, hrHPVs provide a number of potential viral targets for specific HPV antiviral therapies, including the viral encoded oncoproteins E6 and E7, the viral E1 helicase, and the viral E2 regulatory protein.

The hrHPVs associated with cervical cancer encode two oncoproteins, E6 and E7, that are invariably expressed in HPV-positive cancers and target the important cellular growth regulatory proteins p53 and pRb, respectively (1). These cancers are dependent on the continued expression of E6 and E7 and silencing these oncoproteins results in apoptosis and senescence (3, 4). The E6 proteins of the hrHPV types promote the ubiquitin-dependent degradation of the p53 protein (5) through its binding to the E3 ubiquitin ligase E6AP (also known as UBE3A), which ubiquitylates p53 (6–8). In promoting the ubiquitin-dependent proteolysis of p53, E6 counters E7-mediated replication stress signals that activate and stabilize p53 (9). Thus, inhibition of the E6/E6AP-mediated ubiquitylation of p53 is pro-apoptotic and provides a validated target for HPV-positive cancers. For example, direct targeting of E6 induces apoptosis in HPV-positive cancer cells (10). Although the hrHPV E6 proteins have other cellular activities that have been described in the literature, we have focused on the E6/E6AP pathway as a potential therapeutic target for HPV-positive cancers, as well as precancerous lesions.

MicroRNAs (miRNAs/miRs) are small non-coding RNAs (approximately 22 nucleotides in length) that significantly impact and regulate many essential biological pathways. miRNAs negatively regulate gene expression either by translational repression or cleavage of target mRNAs. Many miRNAs have been shown to play roles in cancer, sometimes as oncogenes and other times as tumor suppressors, making them suitable as therapeutic targets. miRNAs themselves can be used as cancer therapeutics as they may affect one or more pathways essential for the replication and survival of specific cancer cells. There have been a number of studies focused on HPV and miRNAs in recent years. These studies have examined whether HPVs, like many other viruses, encode viral miRNAs (11) and whether the HPV oncoproteins regulate the expression of cellular miRNAs (12). Other studies have proposed miRNAs as potential biomarkers of hrHPV infections (13). There have also been studies that examined the effects of miRNA expression in various cancers including cervical cancers (For review see (14)). It was with this mindset that we conducted a screen to identify miRNAs whose transfection into HPV-positive cancer cells stabilize p53.

Here we present a high throughput screen using a cell-based platform to assay p53 stability in hrHPV E6-expressing HeLa cells that identified miRNAs whose transfection increased p53 protein levels. Several of these miRNAs caused apoptosis and/or p21 induction when transfected into hrHPV-positive cells. The goal of this work was to find miRNAs that could be used as therapeutic tools for the treatment of hrHPV-associated diseases, regardless of whether these miRNAs are physiological regulators of p53 stability. To our knowledge, this is the first study to systematically exam miRNAs for their abilities to stabilize p53 in hrHPV-positive cancer cells. We also provide insights into the mechanisms by which some of these miRNAs function in stabilizing p53 and promoting apoptosis.

## Results

### A high throughput screen identified transfected miRNAs that stabilize p53 in HeLa cells

To identify miRNAs that stabilize p53 and induce apoptosis when transfected into hrHPV-positive cancer cells, we performed a high throughput screen in HPV18-positive HeLa cells harboring a reporter for p53 protein stability. The reporter was originally designed by the Elledge lab for their “global protein stability system”(15, 16), and expresses a bicistronic mRNA that encodes an EGFP-p53 fusion protein and the red fluorescent protein DsRed as reference (Figure 1A). The stability of the EGFP-p53 fusion protein is determined by the less stable protein, in this case p53, with a half-life less than 30 minutes in HeLa cells due to E6/E6AP-mediated proteolysis (5, 17). To detect changes in p53 stability, we monitored the EGFP/DsRed fluorescence ratio by laser scanning cytometry. In our screen, for the fusion protein we used a p53 gene encoding the R273C mutant, which is also efficiently targeted by E6/E6AP for ubiquitin-dependent proteolysis (18). Because this mutant is defective in DNA binding (19, 20) and probably in the p53 mitochondrial pro-apoptotic pathways (21), it does not add an extra pro-apoptotic stimulus to the eventual stabilization of the endogenous p53 during the course of the assay. Furthermore, p53(R273C) can interfere with p53 transcriptionally driven apoptosis (22), thus facilitating the acquisition of data for p53 stabilizers that might otherwise kill all cells. The results of this screen, which assayed 885 miRNA mimics, are shown in Figure 1B and Table S1. The top hit, 375-3p, has been previously reported to stabilize p53 in HPV-positive cells (23) providing validation for this screen. Of note, since all the miRNAs used in this study are human, we omitted the use of the prefix hsa-miR in the miRNAs names. We selected for further testing the miRNAs that produced the next 32 highest EGFP/DsRed fluorescence ratios (Table 1).

**Figure 1.**
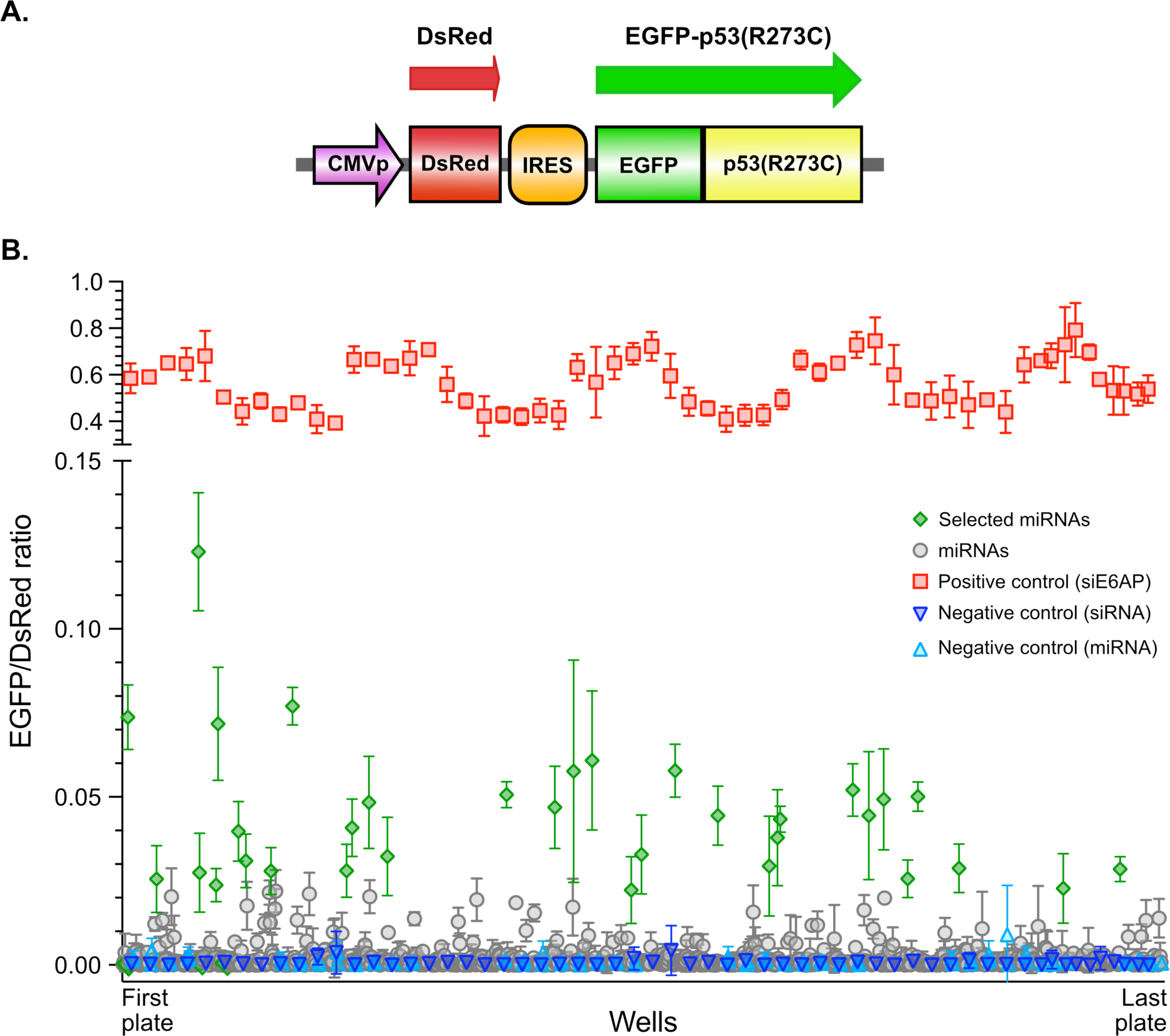
Primary screen. **A.** Reporter cassette of pHAGE-P CMVt RIG2 p53(R273C). The expressed open reading frames are shown with the red and green arrows. CMVp: CMV promoter, IRES: Encephalomyocarditis virus internal ribosomal entry site. **B.** Dot plot showing the effect of the different miRNAs on individual experimental wells during the primary screen. Bars indicate the means of triplicate experiments and the error bars show one standard deviation. The split Y axis has two different scales to help with the visualization of the data.

**Table 1.**
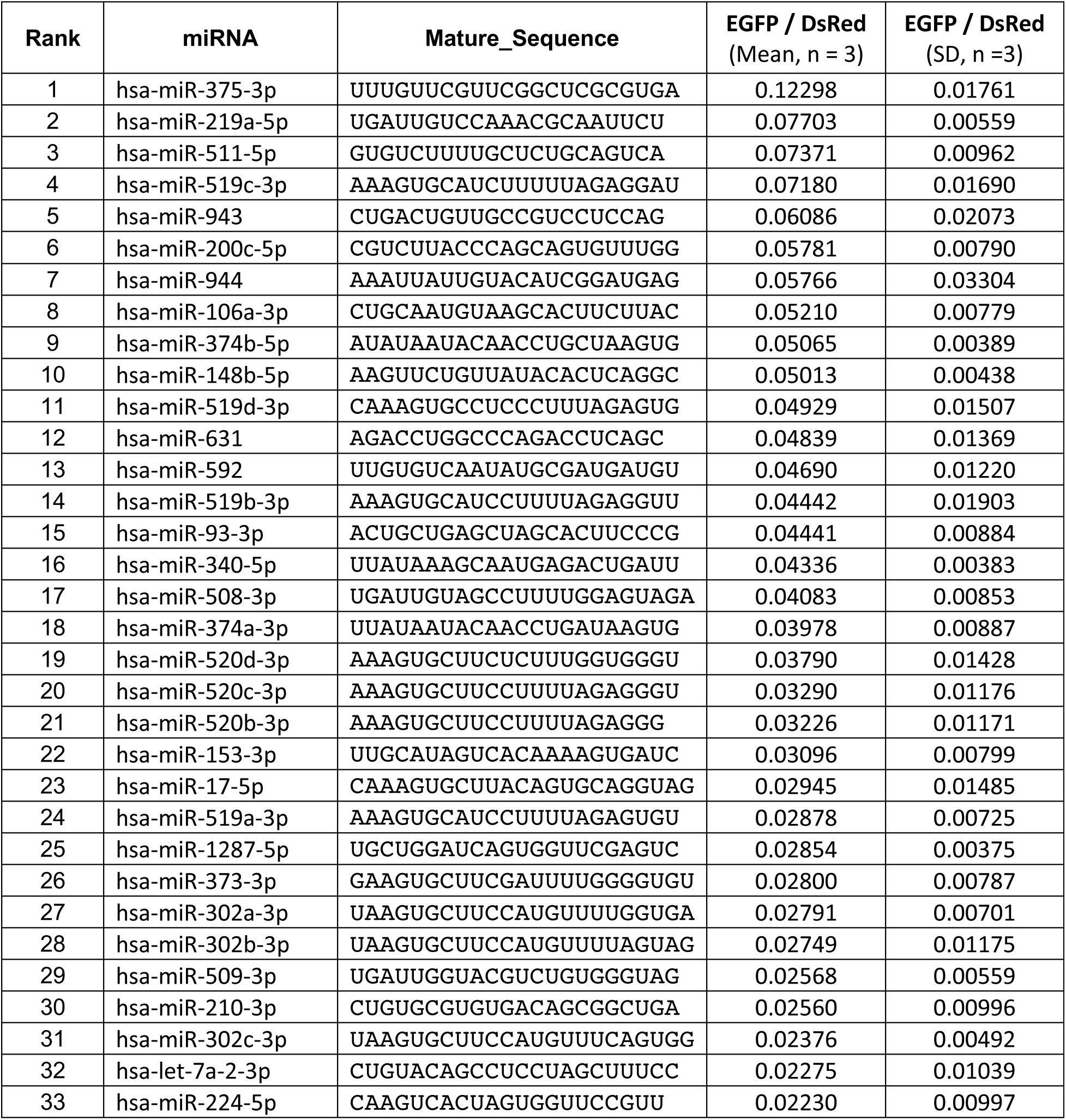
miRNAs selected from primary screen.

To confirm the results obtained in the screen, we tested the selected miRNAs in HeLa cells containing a different bicistronic reporter vector that has been successfully used in a small molecule screen to find p53 stabilizers in HeLa cells (24). In this vector, p53(R273C) is fused to mRuby, a bright monomeric red fluorescent protein, and the reference monomeric green fluorescent protein SGFP2 is fused to H2B (Figure 2A). For this experiment, we determined the percentage of H2B-SGFP2 expressing cells that also expressed mRuby-p53 (R273C) using flow cytometry. Two randomly selected miRNAs (498-5p and 665), which were negative in the initial screen, and the miRNA Ctrl. 1 were used as negative controls, along with 375-3p used as a positive control. As shown in Figure 2B, 25 of the 32 originally selected miRNAs were confirmed as significantly stabilizing mRuby-p53 (R273C) in this reporter HeLa cell line.

**Figure 2.**
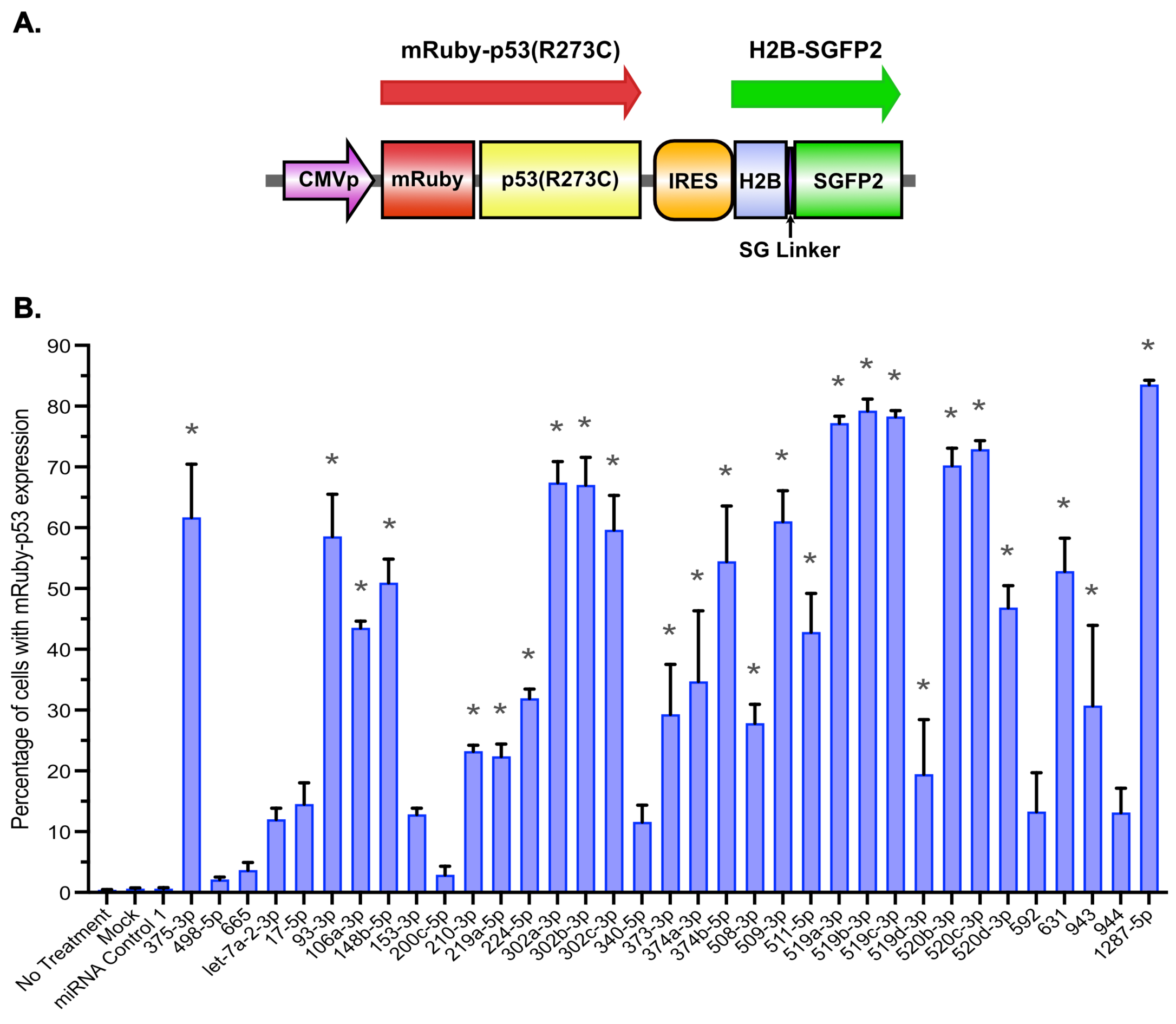
p53 stabilization in HeLa cells expressing the reporter pHAGE-P CMVt RIG3 p53(R273C). **A.** Reporter cassette in pHAGE-P CMVt RIG3 p53(R273C) (24). The expressed fusion proteins are indicated with the red and green arrows. CMVp: CMV promoter, IRES: Encephalomyocarditis virus internal ribosomal entry site, H2B: histone 2B (H2BC11), SG linker: serine-glycine linker. **B.** 72 h after transfection cells were harvested and the percentage of cells expressing mRuby-p53(R273C) was determined by flow cytometry. Columns indicate the means of three independent experiments and the error bars show one standard deviation. Variability between the means was tested using the one-way ANOVA test (F(40, 82) = 84.19 p<0.0001). Then, each sample was compared to non-treated cells using the "Dunnett’s multiple comparisons test" (* = p<0.01).

### Validation of candidate miRNAs in HeLa cells

Twenty seven miRNAs were next tested for their ability to stabilize endogenous p53 in HeLa cells by western blotting (Figure 3 and Table S2), including 3 miRNAs, 17-5p, 153-3p, and 592 that produced a moderate increase in the levels of the reporter p53 . While transfection of most of the miRNAs resulted in some degree of p53 stabilization compared to the mock and negative controls, four miRNAs (219-5p, 106a-3p, 592, and 508-3p) had little or no effect on endogenous p53 levels. The other miRNAs ranged from those that produced a mild to moderate increase in p53 levels (511-5p, 519d-3p, 631, 374a-3p, 153-3p, 17-5p, 373-3p, 210-3p, and 302c-3p) to those that caused stronger p53 stabilization (519c-3p, 943, 374b-5p, 148b-5p, 519b-3p, 93-3p, 520d-3p, 520c-3p, 520b-3p, 519a-3p, 302a-3p, 302b-3p, 509-3p, and 1287-5p). We also examined the effects of the miRNAs on E6AP and HPV18 E6 protein levels (Figure 4A-F and Table S2). Although most miRNAs had little or no effect on E6AP levels, miRNAs 148b-5p, 374a-3p, and 374b-5p decreased the levels of E6AP to values comparable to those observed in cells transfected with the positive control 375-3p, which has been previously shown to target the E6AP mRNA (23).

**Figure 3.**
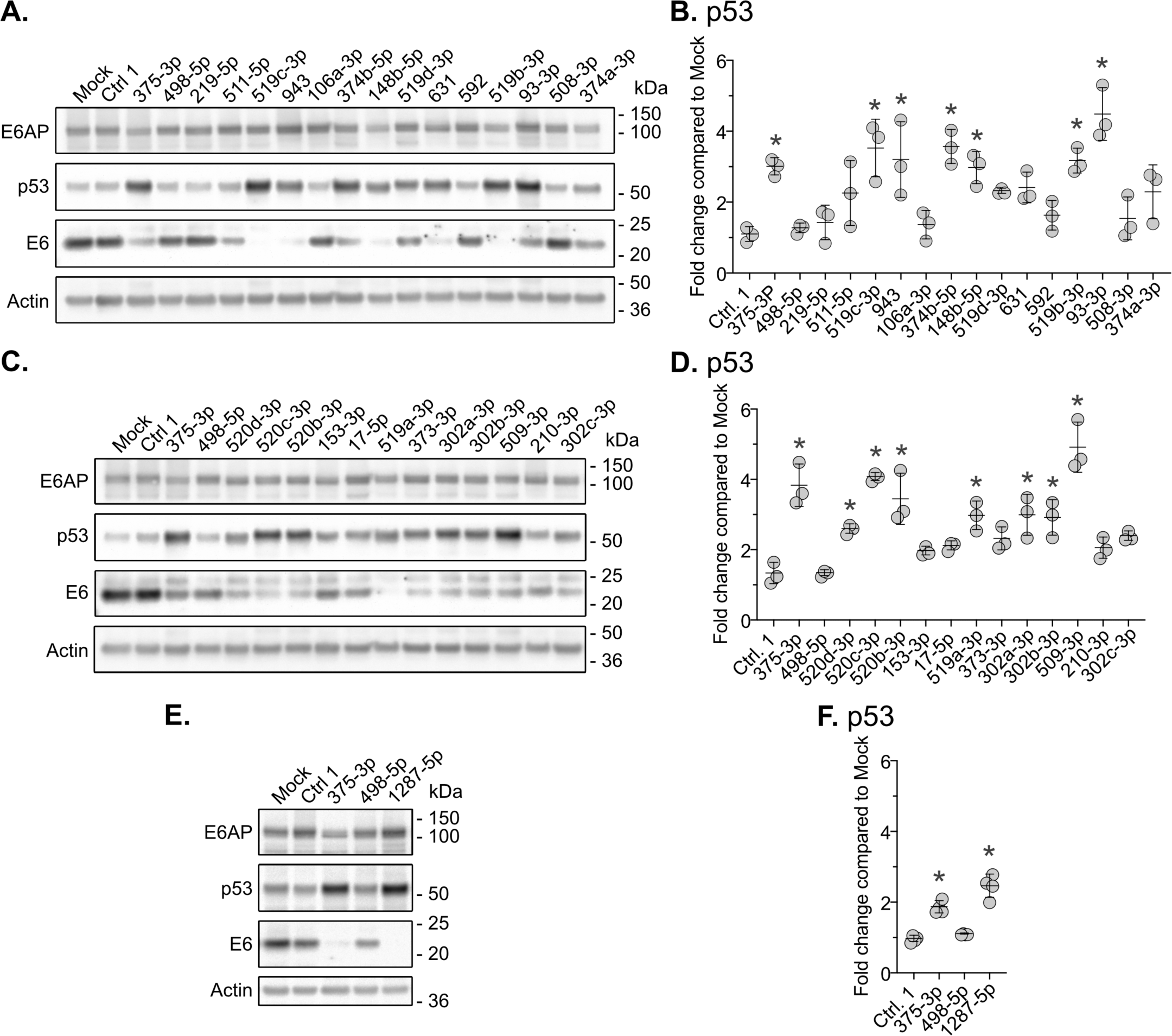
Effect of miRNAs transfection on p53, E6AP, and E6 protein levels in HeLa cells. Cells were lysed 72 h post-transfection and the lysates were resolved by SDS-PAGE. Protein levels of p53, E6AP, E6, and actin were visualized by western blotting. **A, C, and E.** Western blot results from a single experiment. **B, D, and F.** Quantification of p53 from western blot experiments. Values obtained for p53 were normalized to the amount of actin in the same sample and then all values were divided by the corresponding value obtained with the mock transfection in the same blot. Circles represent values from independent experiments and the bars show the mean ± standard deviation. To determine significant differences between the samples in an experiment we used the “one-way ANOVA” test (A. F(16, 34)=8.271 p<0.0001, n=3; B. F(14, 30)=17.76 p<0.0001, n=3; C. F(3, 12)=52.63 p<0.0001, n=4). We then compared each sample with the cells transfected with the Ctrl.1 miRNA using the "Dunnett’s multiple comparisons test" (* = p<0.01).

**Figure 4.**
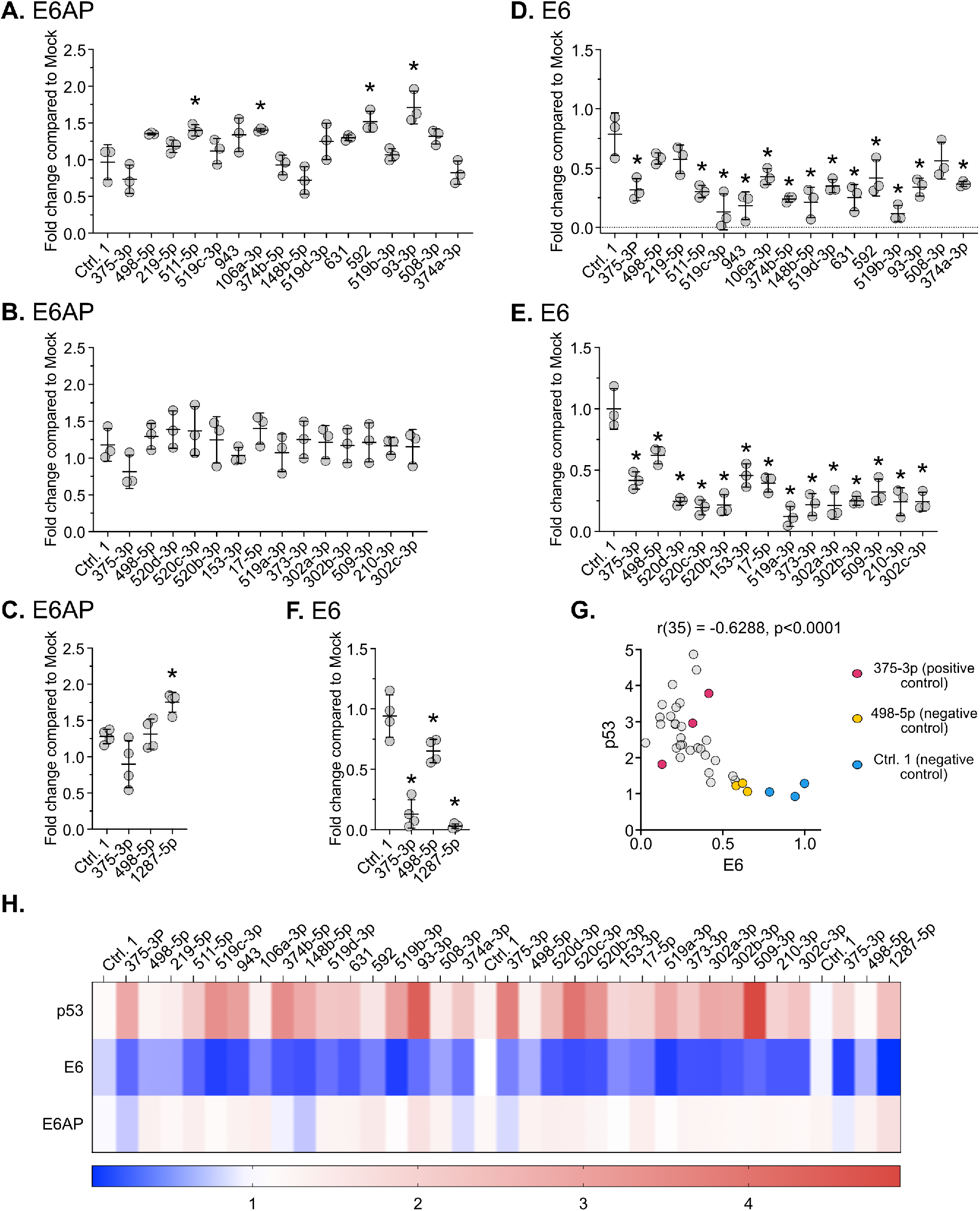
Quantification of the effect of miRNAs transfection on E6AP and E6 protein levels in HeLa cells. Quantification of E6AP (**A-C**) and E6 (**D-F**) protein levels from western blot experiments. The values for E6AP and E6 were normalized to the amount of actin in the same sample, then all values were divided by the corresponding value of the mock transfection in the same blot. Circles show the values of independent experiments and the bars indicate the mean ± standard deviation. The significance of differences between the samples means was determined using the one-way ANOVA test (A. E6AP protein levels F(16, 34)=9.540 p<0.0001, n=3; B. E6AP protein levels F(14, 30)=1.218 p<0.3135, n=3; C. E6AP protein levels F(3, 12)=11.16 p<0.0009, n=4 D. E6 protein levels F(16, 34)=8.672 p<0.0001, n=3; E. E6 protein levels F(14, 30)=18.15 p<0.0001, n=3; F. E6 protein levels F(3, 12)=55.35 p<0.0001, n=4). Each sample was then compared to the cells transfected with the Ctrl.1 miRNA using the "Dunnett’s multiple comparisons test" (* = p<0.01). **G.** The protein levels of p53, and E6 calculated from the western blot experiments were used to analyze whether there is a significant correlation between the levels of these proteins after transfection of HeLa cells with the different miRNAs. This was done by calculating the Pearson’s correlation coefficient (r) and a two tailed p value. **H.** Heat map summarizing the effect of the miRNAs on the protein levels of E6, E6AP, and p53 relative to the cells transfected with Ctrl.1 in the corresponding gel after averaging the quantifications of the western blot experiments showed in Figures 3 and 4, and Table S2).

Transfection of a few miRNAs, most notably 93-3p and 1287-5p, resulted in elevated levels of E6AP. In contrast, transfection of most of the miRNAs tested in this experiment led to a reduction in E6 protein levels. While there was no correlation between the protein levels of E6 and E6AP (r(35) = 0.02098, p = 0.9019) or between E6AP and p53 (r(35) = -0.07389, p = 0.6638), there was a significant negative correlation between the levels of E6 and p53 (r(35) = -0.6288, p<0.0001) (Figure 4G), suggesting that a decrease in E6 expression plays a significant role in the stabilization of p53 by several of these miRNAs in HeLa cells. Examination of these data suggest that, under the conditions used in this experiment, a ∼50% reduction in E6 protein levels appears necessary for p53 stabilization in HeLa cells. Sequence analysis of the validated miRNAs showed a significant enrichment of miRNAs sharing a common seed sequence (5’-AAGUGC-3’) that defines the 302-3p/372-3p/373-3p/519-3p/520-3p family of miRNAs (from now on referred as 302/519 miRNA family) (Figure 5). Based on these results, we selected miRNAs 148b, 374a, 374b, 1287, and five members of the 302/519 family for further analysis.

**Figure 5.**
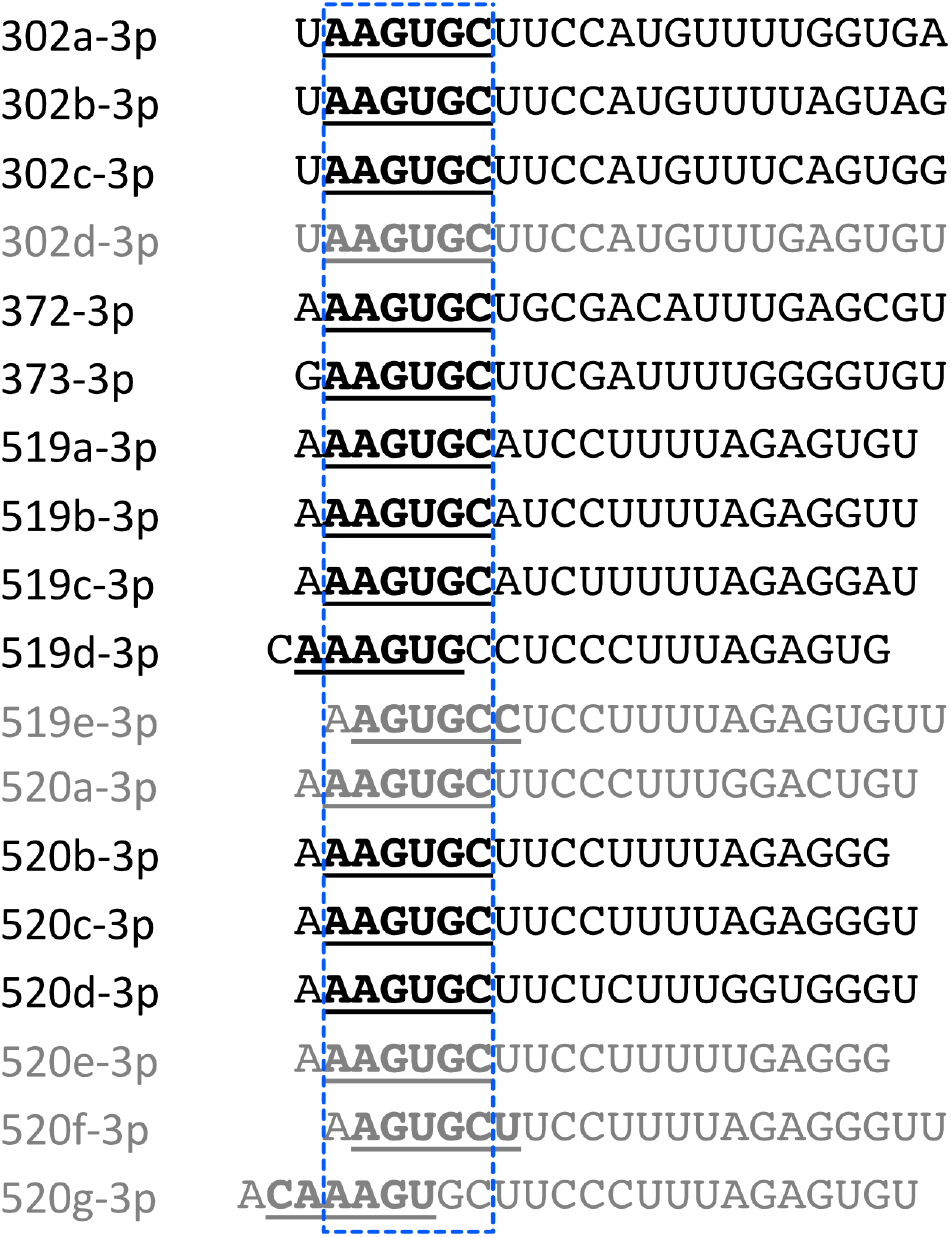
Alignment of members of the 302/519 family of miRNAs. Several of the miRNAs identified in our screen have the same seed sequence indicated with bold letters and underlined. The sequences of the miRNAs were aligned based on the most common seed sequence found in this family framed with a dashed blue line. miRNAs of this family present between the 32 miRNAs selected from the primary screen are indicated with black letters.

### Transfection of selected miRNAs increase p53 in different hrHPV-positive cell lines

To examine whether there was any HPV specificity to the stabilization of p53 by these miRNAs, we examined their effects on the stability of p53 in other hrHPV-positive cancer cell lines. We transfected three cervical cancer cell lines, SiHa (HPV16), ME-180 (HPV68), and MS751 (HPV45), as well as two head and neck squamous carcinoma cell lines, UM-SCC-47 and UM-SCC-104 (both HPV16), with the selected miRNAs and examined p53, E6AP, and actin protein levels by western blot. As shown in Figure 6A-E, the transfection of these miRNAs increased the protein levels of p53 in all HPV-positive cell lines, indicating this effect is conserved in other HPV-associated cervical cancer cell lines independent of the hrHPV type. However, the relative magnitude of the p53 stabilization by individual miRNAs varied in the five HPV-positive cell lines, likely reflecting differences between the cell lines and/or HPV types. 519d-3p consistently produced the lowest increase in p53 protein levels compared to the other 302/519 miRNA family members used in these experiments. These miRNAs share considerable sequence homology throughout and the same seed sequence with exception of 519d-3p, whose seed sequence is displaced one position by the addition of a C at its 5’ end (Figure 5). This suggests that a good pairing of the seed sequence of these miRNAs with their target mRNA is important for the ability of the members of 302/519 miRNA family to induce accumulation of p53 when transfected into hrHPV-positive cancer cells.

**Figure 6.**
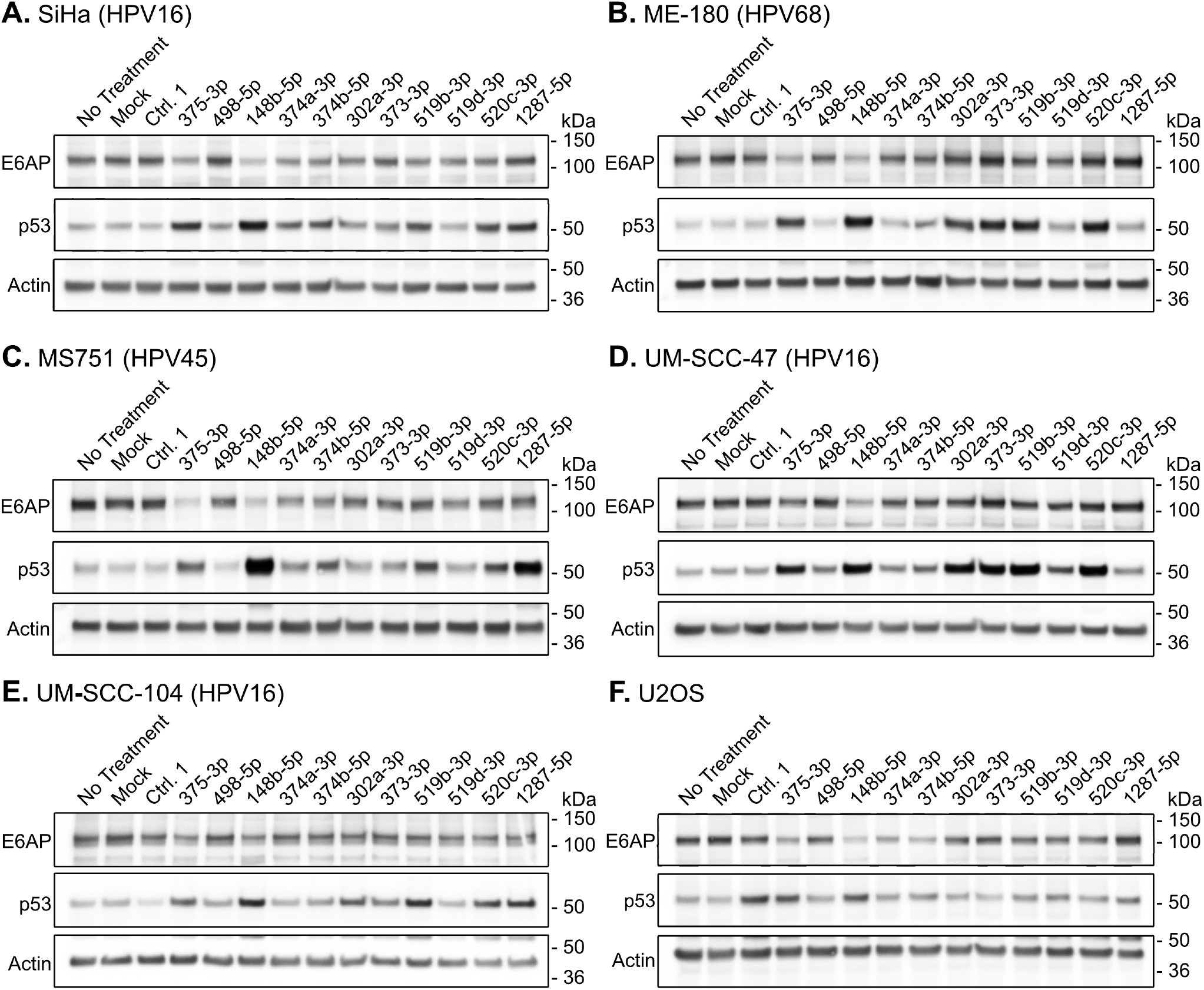
Effect of transfection of selected miRNAs on p53 and E6AP protein levels in different cell lines. 72 h after transfection cells were lysed and the protein extracts were resolved by SDS-PAGE, and the p53, E6AP, and actin proteins levels were determined by western blotting.

To evaluate the involvement of the E6/E6AP pathway in the stabilization of p53 by these miRNAs, we examined their effects on the levels of p53 and E6AP in HPV-negative human U2OS cells (Figure 6F), an osteosarcoma-derived cell line that harbors wt p53 under the control of Mdm2. Transfection of miRNAs 375-3p and 148b-5p increased the levels of p53 in U2OS cells, indicating that these two miRNAs can affect p53 protein levels independent of HPV oncoprotein expression. In contrast, transfection of the miRNAs from the 302/519 family and 1287-5p had minimal or no effect on the levels of p53 in U2OS cells, suggesting that the effect of these miRNAs on p53 stability is dependent upon the expression of hrHPV E6. Of note, transfection of miRNAs 375-3p, 148b-5p and to a lesser degree 374a-3p and 374b-5p, decreased the levels of E6AP in each of the cell lines analyzed.

### Several miRNAs decrease expression of E6/E7

Since the increase of p53 in HeLa cells transfected with the miRNAs identified in this study negatively correlated to the levels of E6 (Figure 4G), we hypothesized that the ability of these miRNAs to stabilize p53 in HPV-positive cancer cells might be mediated, at least in part, by a negative effect on E6 expression. This could be a consequence of destabilizing the E6 protein or decreasing its expression, either by targeting the E6/E7 mRNAs for degradation, preventing their translation, or decreasing viral transcription. We observed that transfection of these miRNAs into HeLa cells, particularly members of the 302/519 family and 1287-5p, decreased both E6 and E7 protein levels similarly (Figure 7A). This is consistent with the idea that these miRNAs affect the expression of the viral E6/E7 mRNAs since the hrHPV E6 and E7 genes are expressed from a single promoter located in the viral LCR. This effect on the level of the E6/E7 mRNA in HeLa cells was confirmed by qRT-PCR using primers that detect all HPV E6/E7 transcripts (Figure 7B). Interestingly, all miRNAs tested, including 375-3p and 148b-5p that increased p53 protein levels in the absence of hrHPV proteins (Figure 6F), also decreased the levels of the E6/E7 transcripts. Although 498-5p reduced the level of the E6/E7 mRNAs in HeLa cells, consistent with the reduced level of E6 protein shown in Figures 3, 4 and 7A, we did not observe an increase in p53 protein levels (Figures 3 and 6). While 375-3p was as efficient in repressing the expression of the E6/E7 mRNAs in HPV16 SiHa cells as in HPV18 HeLa cells, most of the tested miRNAs were less efficient decreasing the levels of the E6/E7 transcripts in SiHa cells, even though they were efficiently transfected as determined by siGlo RISC-free transfection control (data not shown). In addition to 375-3p, only 519b-3p and 1287-5p showed comparable effects on the E6/E7 transcripts in both HPV-positive cell lines. Surprisingly, 148b-5p, which stabilized p53 in SiHa cells, produced an increase in the levels of the E6/E7 mRNAs but we could not confirm whether E6 protein levels were affected in this cell line (e.g. by preventing translation) because we were unable to detect HPV16 E6 using any of the commercially available anti-HPV16 E6 antibodies.

**Figure 7.**
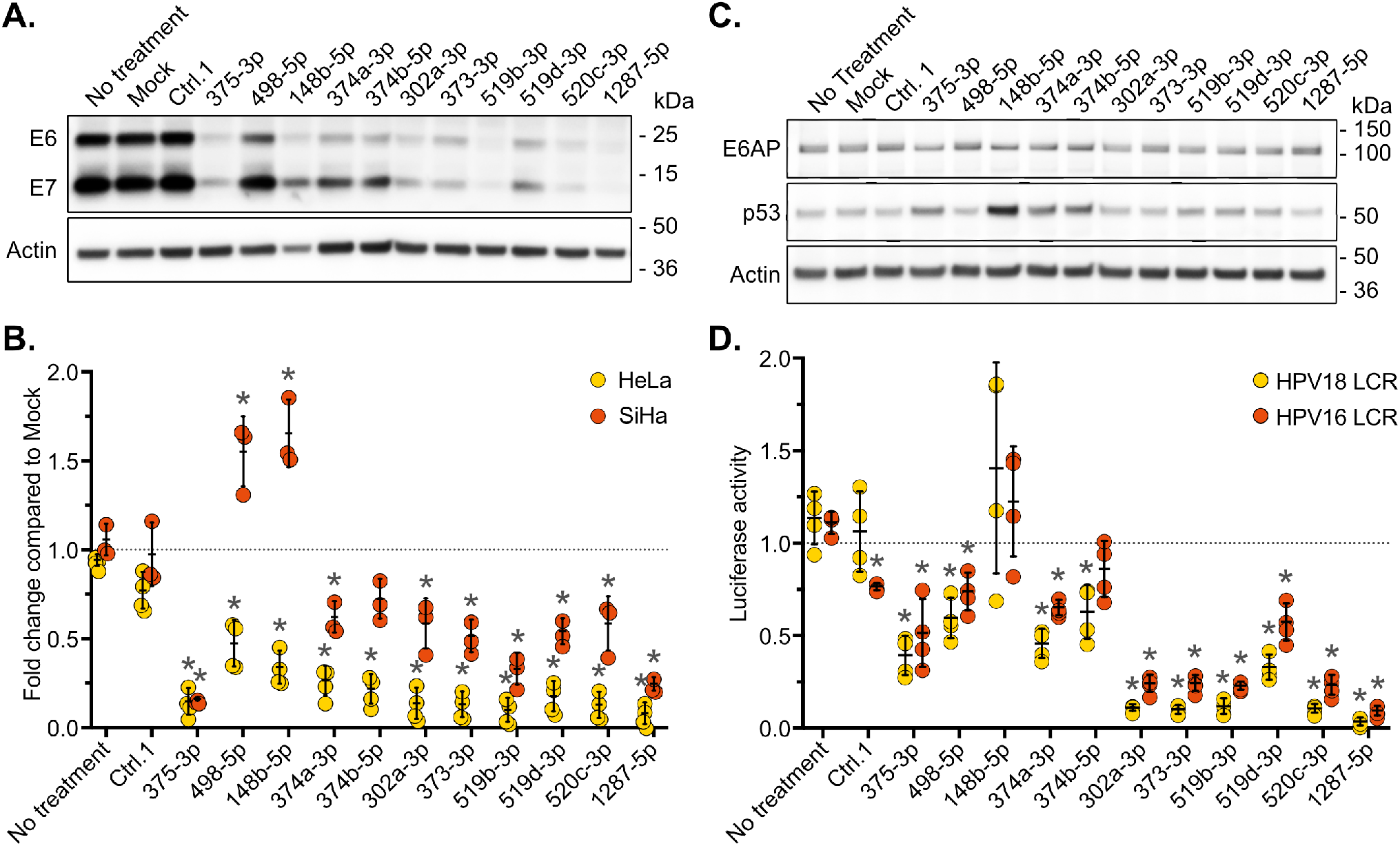
Effect of selected miRNAs on HPV E6 and E7 expression. Cells were harvested 72 h post-transfection and then used for the following experiments. **A.** Western blot showing the effect of the selected miRNAs on E6, E7, and actin protein levels in HeLa cells. **B.** Expression of the E6/E7 mRNAs after transfection of the selected miRNAs was quantitated by QRT-PCR. Measures from each experiment were normalized to the value of the corresponding mock transfection. Differences between samples were analyzed by the “one-way ANOVA” test (HeLa F(12, 39) = 42.32 p<0.0001, n=4; SiHa F(12, 26) = 41.38 p<0.0001, n=3), then each sample was compared to the corresponding non-treated cells using the "Dunnett’s multiple comparisons test" (* = p<0.01). **C.** Western blot showing the effect of transfecting the indicated miRNAs into 1321 cells on the p53, E6AP, and actin proteins levels after 72 h incubation. **D.** Effect of transfection of selected miRNAs on the promoter activity of the HPV16 and HPV18 LCRs using a luciferase reporter assay in C33A cells. Values from each experiment were normalized to the value obtained for the corresponding mock transfection. Differences between samples were analyzed by the “one-way ANOVA” test (HPV 16 LCR, F(12, 38) = 34.91 p<0.0001, n=4; HPV 18 LCR, F(12, 39) = 23.47 p<0.0001, n=4), then each sample was compared to the corresponding non-treated cells using the "Dunnett’s multiple comparisons test" (* = p<0.01). In B and D, circles represent values from independent experiments and the bars show the mean ± standard deviation.

To determine whether these miRNAs were targeting the untranslated regions (UTR) of the HPV16 E6/E7 mRNAs, we examined their effect in 1321 cells, a human keratinocyte cell line that expresses HPV16 E6/E7 under the control of the human β-actin promoter and from a mRNA containing the 3’UTR of the human β-actin mRNA (25). In this cell line, the 302/519 miRNA family members (302a-3p, 373-3p, 519b-3p, 519d-3p, and 520c-3p) and 1287-5p failed to increase the p53 protein levels, suggesting that their effect involves actions over the promoter activity of the HPV16 LCR and/or the 3’UTR of the E6/E7 transcripts (Figure 7C). In contrast, 375-3p, 148b-5p, 374a-3p, and 374b-5p transfection resulted in an increase in p53 protein levels, indicating it is likely through a different mechanism than the miRNAs of the 302/519 family and 1287-5p. Considering that the E6/E7 transcripts in the different hrHPV-positive cell lines tested have different 3’UTRs, depending on the viral sequence inserted in the host cell and the insertion site, it is more likely that the miRNAs of the 302/519 family and 1287-5p affect E6 and E7 protein levels by acting on the viral LCR promoter activity. In line with this hypothesis, 1287-5p and the 302/519 family members efficiently repressed the HPV16 LCR and HPV18 LCR promoter activity in a reporter system in the HPV-negative cervical cancer cells C33A (Figure 7D),

### Selected miRNAs induce apoptosis and/or p21 expression

Stabilization of p53 is expected to be pro-apoptotic in hrHPV-positive cells. We therefore tested whether miRNAs that stabilized p53 in HeLa cells induce apoptosis. Flow cytometric analysis of HeLa cells transfected with different miRNAs and stained with the apoptosis marker Annexin V revealed an increase in the fraction of apoptotic cells when compared to untreated or mock transfected cells (Figure 8A). Of note, miRNAs Ctrl.1 and 498-5p, which did not increase p53 levels in hrHPV-positive cells (Figures 3 and 6), did increase Annexin V staining upon transfection into HeLa cells. In contrast, there was a correlation between induction of apoptosis and stabilization of p53 in HeLa cells transfected with the 302/519 family members (Figure 3). In this family, 519d-3p, which produced the lowest increase in p53 levels in HeLa cells, was also the least efficient in inducing apoptosis.

**Figure 8.**
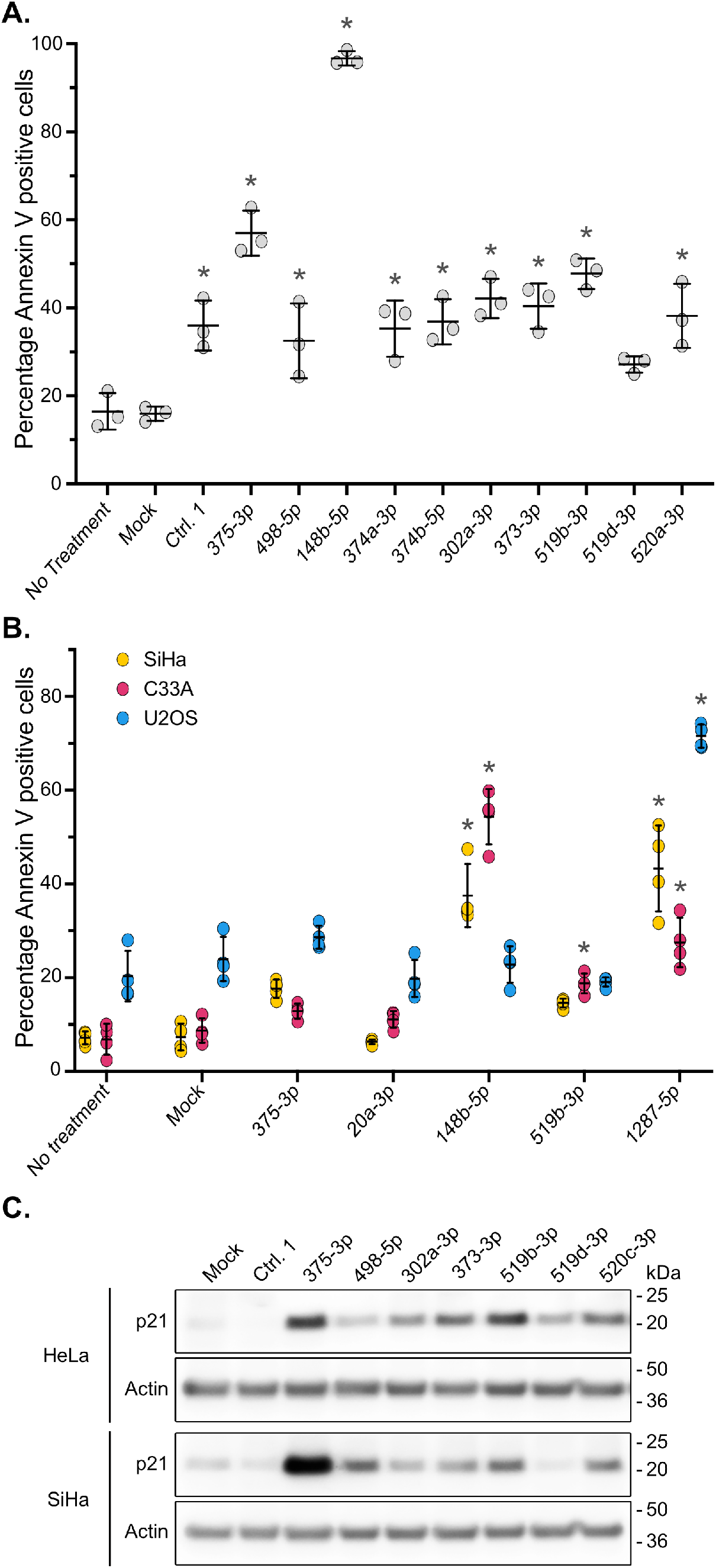
Transfection of selected miRNAs induces apoptosis in HeLa cells. **A.** Cells were harvested 72 h post-transfection and stained with Annexin V as an indicator of apoptosis. The percentage of cells stained with Annexin V was determined by flow cytometry. Variability between samples was first analyzed with the one-way ANOVA test (F(12,26)=48.04 p<0.0001, n=3). Then each sample was compared to the non-treated cells with the "Dunnett’s multiple comparisons test" (* = p<0.01). Circles represent values from independent experiments and the bars show the mean ± standard deviation. **B.** Cells were transfected with the indicated miRNAs and 72 h later the cells were harvested and stained with Annexin V as a marker for apoptosis. The percentage of Annexin V positive cells was quantitated by flow cytometry. Variability between samples for each cell line was analyzed with the one-way ANOVA test (SiHa F(6,21)=45.17 p<0.0001, n=4; C33A F(6,21)=85.96 p<0.0001, n=3; U2OS F(6,21)=104.07 p<0.0001, n=4). Each sample was then compared to the non-treated cells with the "Dunnett’s multiple comparisons test" (* = p<0.01). Circles represent values from independent experiments and the bars show the mean ± standard deviation. **C.** Effect of the indicated miRNAs on the p21 and actin proteins levels HeLa and SiHa cells visualized by western blotting 72 h post transfection.

We next examined the ability of a subset of these miRNAs to induce apoptosis in SiHa cells and two HPV-negative cancer cell lines, C33A and U2OS, using Annexin V staining (Figure 8B). Transfection with the miRNA 20a-3p was utilized as a negative control, since it does not induce apoptosis in cervical cancer cells (26). The 1287-5p miRNA was also included in these experiments as it was found to stabilize p53 (Figures 3 and 6) and to efficiently decrease E6/E7 mRNAs expression in both HeLa and SiHa cells (Figure 7B). Similar to their effect in HeLa cells, 148b-5p and 1287-5p induced apoptosis in SiHa cells whereas 375-3p and 519b-3p had very little or no effect. While all of these miRNAs increased the levels of p53 when transfected into SiHa cells (Figure 6A), 148b-5p and 1287-5p led to the highest p53 levels. Both 148b-5p and 1287-5p stimulated apoptosis in C33A cells, an HPV-negative cervical cancer cell line that already expresses high levels of an inactive mutant form of p53 (27), indicating that these miRNAs can induce apoptosis independently of p53.

Since 519b-3p has been reported to induce senescence in HeLa cells through p21 expression, which is partially dependent of p53 (28), we tested whether other members of the 302/519 family have a similar effect on p21 protein levels and if it also occurs in HPV16-positive SiHa cells. Since 375-3p has been shown to increase p21 protein levels in both HeLa and SiHa cells (23), it was included as the positive control. As shown in Figure 8C, the miRNAs of the 302/519 family tested in this experiment increased p21 protein levels in both cell lines with the exception of 519d-3p, which did not increase p21 levels in SiHa cells. Interestingly, 498-5p led to an increase in p21 protein levels in SiHa cells; however this is likely to be at least partially p53 independent since this miRNA failed to induce a marked p53 accumulation under similar conditions in these cells (Figure 6A).

### miRNAs of the 302/519 family and 1287-5p induce p53-dependent apoptosis in HeLa cells

We next examined if induction of apoptosis by several representative miRNAs is p53-dependent in HeLa cells. We performed Annexin V staining of HeLa cells co-transfected with the miRNAs and an siRNA targeting p53 or its C911 variant (29) to address possible off-target effects of the siRNA (Figure 9A). As expected, the positive control 375-3p increased the percentage of Annexin V-stained cells while 20a-3p had no significant effect when compared to untreated cells. Surprisingly, the apoptotic effect of 375-3p was independent of p53 since knock-down of p53 had little effect on the percentage of apoptotic cells in HeLa cells co-transfected with this miRNA. The effect of 498-5p on apoptosis is also p53-independent and consistent with the inability of this miRNA to stabilize p53 when transfected into hrHPV-positive cell lines (Figures 3 and 6). Similarly, the apoptotic effects of miRNAs 148b-5p and 374a-3p are also independent of p53, as co-transfection of these miRNAs with either the siRNA against p53 or its C911 version have similar effects on the percentage of apoptotic cells. This indicates that, in these cases, the effect observed when p53 is knocked down can be attributed to off-target effects of the siRNA against p53. In contrast, the numbers of apoptotic cells observed with 519b-3p (as a representative of the 302/519 family of miRNAs) and 1287-5p were reduced when co-transfected with the p53 siRNA but not with its C911 version, suggesting that the apoptotic effect of these miRNAs is largely dependent on p53 in HeLa cells. Similar results were observed when using PARP cleavage as a marker for apoptosis in western blot analysis (Fig 9B).

**Figure 9.**
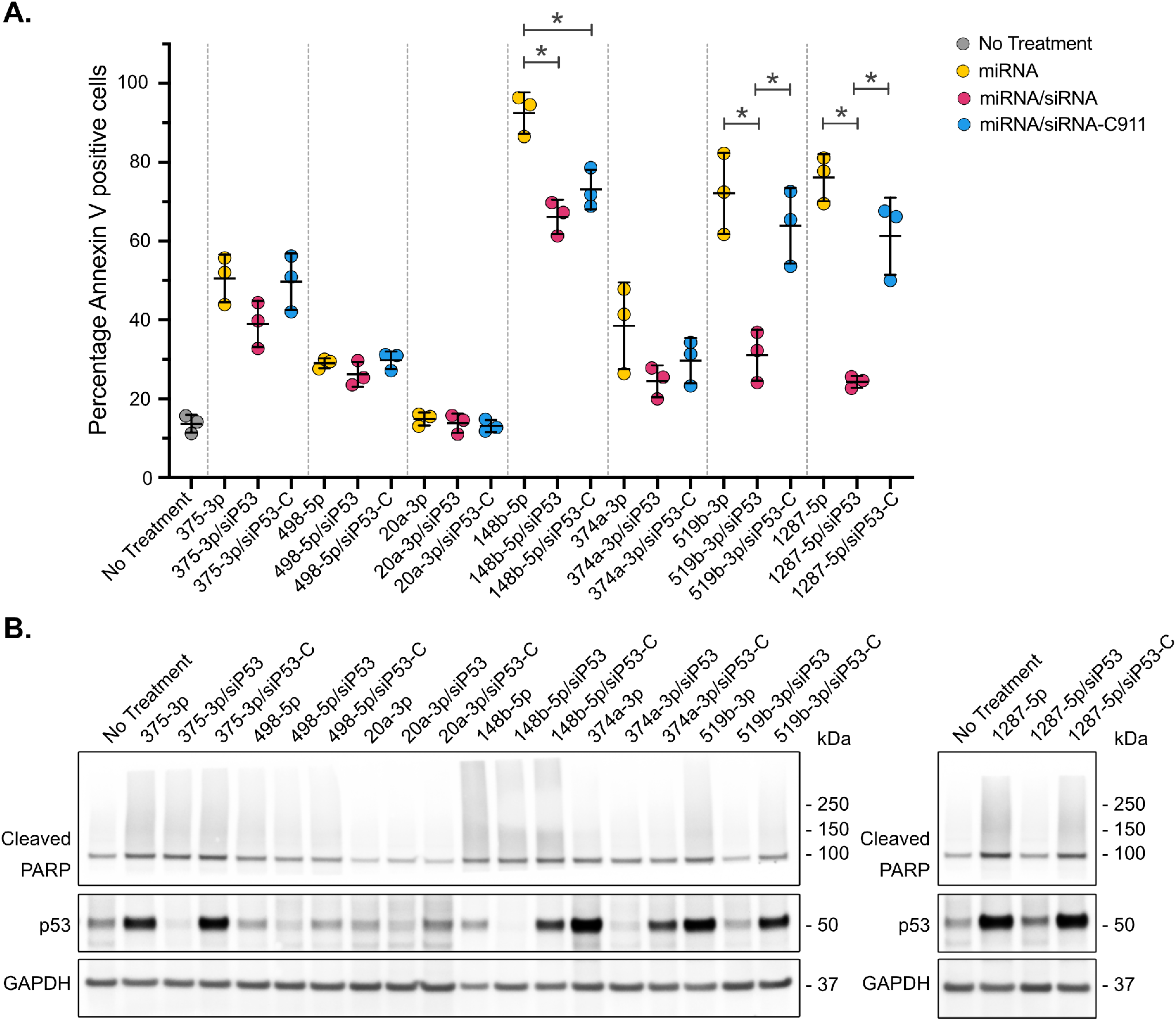
p53 dependence of apoptotic effect of selected miRNAs in HeLa cells. **A.** Cells were transfected with the indicated miRNAs either alone or together with a siRNA against p53 or its C911 variant. After incubation for 72 h the cells were harvested and stained with Annexin V as a marker for apoptosis. The percentage of Annexin V stained cells was determined by flow cytometry. The values obtained for the three transfections with each miRNA were first analyzed with the one-way ANOVA test (375-3p F(2, 6) = 3.075 p=0.1204; 498-5p F(2, 6) = 1.914 p=0.2276; 20a-3p F(2, 6) = 0.6553 p=0.5528; 148b-5p F(2, 6) = 23.57 p=0.0014; 374a-3p F(2, 6) = 2.681 p=0.1472; 519b-3p F(2, 6) = 17.71 p=0.0030; 1287-5p F(2, 6) = 48.00 p=0.0002). For all samples n=3. Then, the values from samples transfected with the same miRNA were compared with each other using the "Tukey’s multiple comparisons test" (* = p<0.01). **B.** 72 h post-transfection cells were lysed and the p53, cleaved PARP, and GAPDH proteins levels were determined by SDS-PAGE and western blotting.

## Discussion

In this study we identified several miRNAs that increased p53 protein levels when transfected in HeLa cells. Some of the miRNA hits from our high throughput screen (148b-5p, 302a-3p, 373-3p, and 374b-5p) have been previously reported to have decreased expression in cervical cancer cells and tissues compared to the controls used in each of the studies (26, 30–32). Another positive miRNA from the screen, 1287-5p, has been found to have no change in its expression level in cervical cancer cells, but is sequestered away from its targets by the circular RNA circSLC26A4, which is overexpressed in hrHPV-positive cancer cells (33). Together these findings suggest that interfering with the functions of some of the miRNAs identified in this work might contribute to the establishment of HPV infections or/and progression of hrHPV-infected cells towards cancer. Consistent with that, several of the miRNAs identified in our screen, namely 148b-5p, 302a-3p, 374b-5p, or 519b-3p, have been shown to slow cell growth and induce apoptosis in hrHPV-positive cervical cancer cells (28, 31, 32, 34, 35). Taken together, these previous findings validate this screening platform as a reliable tool to identify miRNAs that affect the cell growth and survival of hrHPV-positive cells, as expected from miRNAs whose transfection stabilize p53.

Since the ubiquitin ligase activity of E6AP/E6 is responsible for the efficient proteolysis of p53 in hrHPV-positive cervical cancer cells, we determined whether transfection of any of the miRNAs identified in this screen altered the levels of E6AP in HeLa cells. Interestingly, only three of the 27 miRNAs analyzed in HeLa cells, 148b-5p, 374a-3p, and 374b-5p, led to a decrease in E6AP protein levels comparable to the decrease produced by 375-3p. Although we did not determine whether they directly bind to the E6AP mRNA, target prediction analysis using the miRDB database (www.mirdb.org) (36, 37) revealed that 148b-5p and 374b-5p each have three sites complementary to their seed sequences within the 3’UTR of the E6AP mRNA. Remarkably, these four miRNAs (148b-5p, 374a-3p, 374b-5p, and 375-3p) also led to decreased E6 expression. This simultaneous decrease of both E6AP and E6 may be required to stabilize p53 in HeLa cells, since a small amount of E6AP is sufficient to efficiently promote p53 degradation in this cell line (38). The need to suppress the very efficient E6/E6AP-mediated degradation of p53 to increase the levels of this protein in hrHPV-positive cancer cells is exemplified by miRNAs 375-3p and 1287-5p. Both miRNAs stabilized p53 in absence of E6 when transfected into U2OS cells, however, they also efficently decrease the levels of E6 when transfected into HeLa and SiHa cells, suggesting that this is a necessary condition to stabilize p53 in these cells. Moreover, in this study we haven’t found any miRNA that stabilize p53 in hrHPV-cancer cells that do not significantly decrease the E6 protein level too. In contrast, transfection of 498-5p, which also resulted in decreased levels of E6 in these cells, did not stabilize p53 in HeLa cells, suggesting that the remaining E6 protein was sufficient to target p53 for degradation. In line with this idea, in our study, miRNAs that failed to reduce E6 protein levels to approximately half of its original amount or beyond also failed to increase p53 levels in HeLa cells, suggesting that there is a considerable excess of E6 in these cells that must be overcome to stabilize p53. High levels of expression of E6 could therefore be a limiting factor for the use of some miRNAs such as 498-5p as therapeutics, at least in some hrHPV-associated cancers.

While several of the transfected miRNAs were found to induce apoptosis, the mechanisms by which this occurred varied. The apoptotic induction in HeLa cells for three of the miRNAs, 148b-5p, 374a-3p, and 375-3p, was independent of p53 stabilization. In contrast, transfection of 1287-5p induced apoptosis in HeLa cells in a p53-dependent manner. The miRNA 1287-5p was also found to efficiently induce apoptosis in U2OS cells without a noticeable increase in p53 levels and to increase the number of apoptotic C33A cells, which do not express wt p53, indicating that this miRNA promotes cell death independent of p53 stabilization in these HPV-negative cells. Therefore, 1287-5p can induce apoptosis via different pathways depending on the cell type. This is also exemplified by Ctrl. 1 whose transfection stabilized p53 only in U2OS cells. The other miRNA that we found to induce apoptosis in HeLa cells in a p53-dependent manner upon transfection was 519b-3p, a member of the 302/519 family. Abdelmohsen and colleagues have previously reported that 519-3p transfection in HeLa cells increases p21 levels in a partly p53-independent manner, resulting in growth inhibition and cell survival (28). There are a number of technical differences between our study and that of Abdelmohsen et al. which could account for the discordant results, including different miRNA concentrations and end points. However, we both found that 519b-3p transfection did not induce apoptosis in SiHa cells, but led to an increase in p53 and p21 protein levels. It has been reported that p53 must accumulate to a threshold level to induce apoptosis (39). Thus, it is feasible that under the experimental conditions used in our study 519b-3p did not increase p53 levels to the threshold necessary to trigger apoptosis in SiHa cells. In addition, we observed that transfection of all miRNAs belonging to the 302/519 family, with the exception of 519d-3p in SiHa cells, increased p21 protein levels in hrHPV-positive cells indicating this is a family characteristic rather than a feature of 519b-3p alone.

Throughout this study, 519d-3p has been less efficient than the other members of the 302/519 family of miRNAs in promoting increased p53 and p21 protein levels in the hrHPV-positive cell lines, as well as in repressing expression from the HPV16 and HPV18 LCRs, and inducing apoptosis in HeLa cells. In addition to a few base substitutions in the 3’ end sequences, a difference between 519d-3p and the other miRNAs of this family included in this study is the addition of a C at its 5’end. This additional nucleotide produces a displacement of the miRNA sequences, including its seed sequence, which is 5’-AAAGUG-3’ instead of 5’-AAGUGC-3’, the most common seed sequence between the members of the 302/519 family. Since a seed sequence match is very important for miRNA target recognition (40, 41), this change in 519d-3p’s seed sequence is expected to impact its functions when compared to the other members of the 302/519 family. The data presented in this study supports this hypothesis and indicate that the correct matching of the seed sequence 5’-AAGUGC-3’ with their target mRNAs is an important factor mediating the effects of this family of miRNAs on p53 and p21 protein levels as well as on cell growth and survival of hrHPV-positive cancer cells.

Since their discovery in 1993 (42, 43), the interest in miRNAs continues to increase. Research in miRNAs is being translated into diagnostics as miRNAs are considered sensitive and specific biomarkers. In contrast, their use as therapeutics have lagged with only a few clinical trials currently underway (reviewed in (44, 45)). This may be due in part to the lack of single target specificity characteristic of miRNAs. This multiplicity of targets makes miRNAs potentially challenging as therapeutics because of the possibility of unexpected and/or undesirable side effects. However, this property makes them an attractive tool for cancer treatment since miRNAs can simultaneously regulate multiple cellular pathways by modulating the expression of several genes in a coordinated fashion. The utility of miRNAs as therapeutic tools will be improved by a more complete understanding of how they function in the cellular networks in which they are embedded. In this study, we identified several miRNAs that stabilize p53 in hrHPV-cancer cells and induce apoptosis in HeLa cells in a p53-dependent or in p53-independent manner. Despite the heterogenicity of the responses to different miRNAs, decreasing E6 protein levels seems to be required to increase p53 protein levels in HeLa cells since all miRNAs found to stabilize p53 also affected the expression of E6. Further research into the mechanisms involved in the stabilization of p53 and/or induction of apoptosis by these miRNAs in hrHPV-positive cells will improve our understanding of the hrHPV-host interactions and contribute to the identification of new potential therapeutic targets for hrHPV infections and cancers.

## Materials and methods

### Plasmids

To create a lentiviral reporter vector to monitor the protein levels of p53 *in vivo* in hrHPV cancer cells, the p53 open reading frame (ORF) carrying the mutation R273C was first amplified by PCR using the primers hp53-1 (5’-CACCATGGAGGAGCCGCAGTCAGATCC-3’) and hp53-8 (5’-GGATCCTCAGTCTGAGTCAGGCCCTTCTGTCTTG-3’) and cloned into the pENTR/D-TOPO vector (ThermoFisher) generating the entry vector pENTR p53(R273C) (p6140). The p53(R273C) ORF was then recombined from the entry vector into a lentiviral vector containing a GPS (<u>G</u>lobal <u>P</u>rotein <u>S</u>tability) cassette (gift from S. Elledge) using the GATEWAY cloning system (ThermoFisher), resulting in the vector pHAGE-P CMVt N-RIG2 p53(R273C) (p7987). The GPS reporter cassette expresses a bicistronic mRNA encoding EGFP fusion proteins and the red fluorescent protein DsRed as reference (15, 16, 46, 47). The reporter vector pHAGE-N CMVt N-RIG3 p53(R273C) (p7709), which contains a variant of the GPS reporter cassette, expresses p53(R273C) fused to the monomeric red fluorescent protein mRuby and a monomeric green fluorescent protein (SGFP2) fused to a histone 2B (H2BC11) as reference (38).

### Cell lines

HeLa, HeLa RIG2 p53(R273C), HeLa RIG3 p53(R273C), Siha (ATCC® HTB-35™), C33A (ATCC® HTB-31™), U2OS (ATCC® HTB-96™), ME-180 (ATCC® HTB-33™), MS-751 (ATCC® HTB-34™), UM-SCC-47 and UM-SCC-104 (EMD-Millipore), and human keratinocytes immortalized by HPV16 E6 and E7 expressed from the β-actin promoter (p1321) (25) referred to here as 1321 cells were grown in Dulbecco’s modified Eagle’s medium (DMEM) supplemented with 10% (v/v) fetal bovine serum at 37°C 5% CO2.

### miRNA screen

For the primary miRNA screen, HeLa RIG2 p53(R273C) cells were reverse transfected with the Ambion Human Pre-miR miRNA Mimic Library (2009) based on miRBase release 13.0. in a one-target, one-well format using 384-well microtiter plates in a high throughput format. The Z’-factor (48) was calculated for each assay plate and was consistently > 0.6, indicating assay robustness. siRNA buffer (1×) (Horizon #B-002000-UB-100) was aliquoted into wells, miRNA mimics were added so that the final concentration was 40 nM/well, and DharmaFECT1/OptiMEM was dispensed into wells. While the miRNA/lipid mix was allowed to complex, cells were trypsinized, counted, and resuspended to reach a plating density of 600 cells/well. Cells were seeded on top of the miRNA/lipid mixture, briefly centrifuged, and incubated for 72 hours, a time that we determined to be optimal for p53 stabilization for small miRNAs. The cells were then equilibrated to room temperature (∼15 min) before analysis. Using TTP LabTech’s Acumen eX3 laser scanning cytometer, the total EGFP and DsRed intensities in each well were quantitated. Internal controls were present in Columns 13 and 14 of each library plate (Pre-miR miRNA Precursor Molecules Negative Control #1 - Ambion AM17110; Pre-miR miRNA Precursor Molecules Negative Control #2 – Ambion AM17111; PLK1 SMARTpool Dharmacon M-003290-01). In addition, siRNAs against E6AP (UBE3A, Dharmacon D-005137-04) and non-targeting siRNAs (Dharmacon D-001210-01 and D-001210-02) were added to each plate. The miRNA mimics were screened in triplicate. High throughput libraries and screening capability was provided by the ICCB-Longwood Screening Facility at Harvard Medical School.

### miRNA transfections

All cells were reverse transfected with miRNAs (Table 2) using RNAiMax according to manufacturer’s instructions (ThermoFisher Scientific). More specifically, 750 µl of suspended cells were added to 250 µl of a transfection mix containing 2 µl of Lipofectamine RNAiMax and different miRNAs for a final concentration of 10nM and plated in 12-well plates. miRNA transfection efficiency in different cell lines was confirmed using siGlo Red Transfection Indicator (Horizon, D-001630-02-05).

**Table 2.**
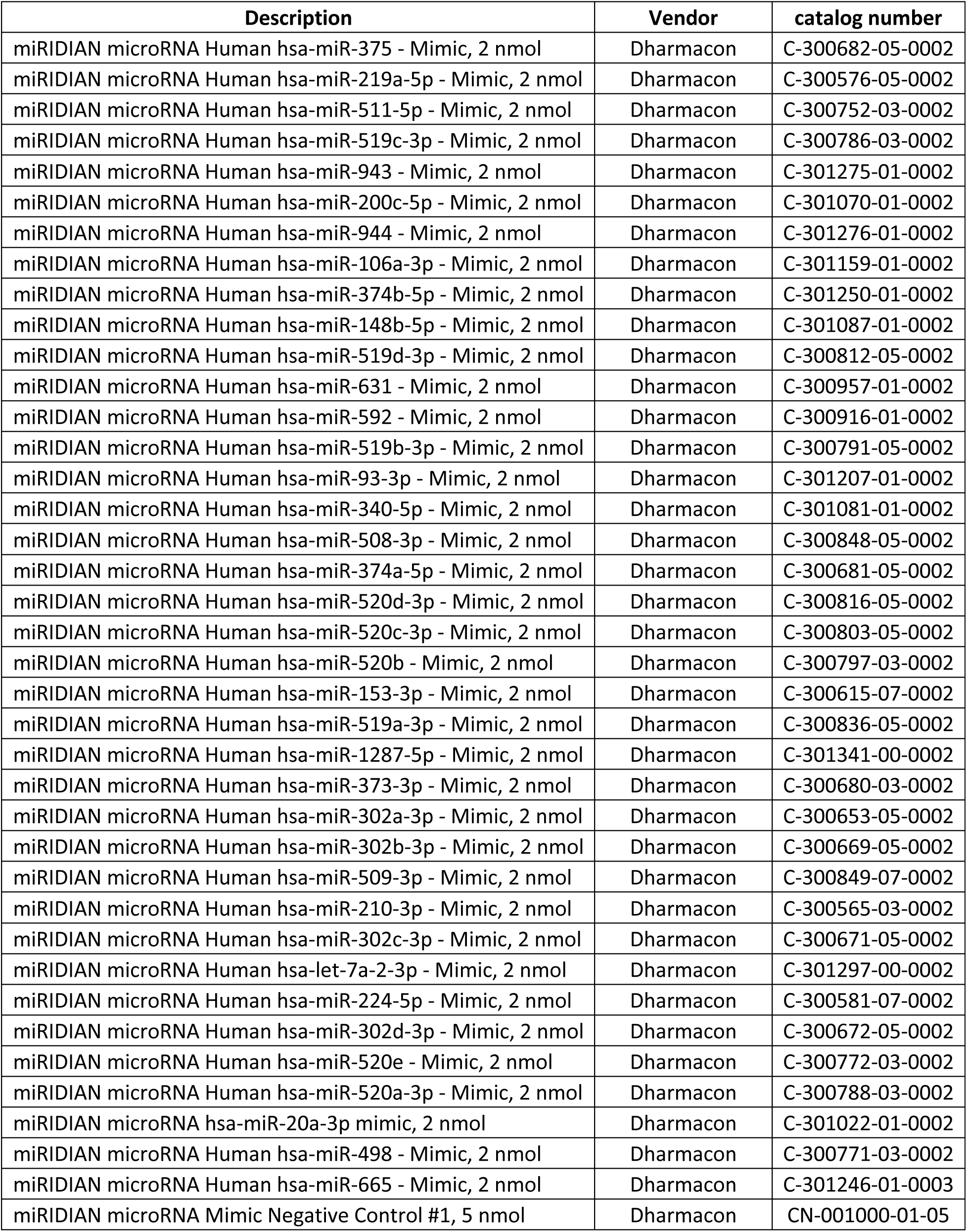
miRNAs used in this study.

| Description | Vendor | catalog number |
| --- | --- | --- |
| miRIDIAN microRNA Human hsa-miR-375 - Mimic, 2 nmol | Dharmacon | C-300682-05-0002 |
| miRIDIAN microRNA Human hsa-miR-219a-5p - Mimic, 2 nmol | Dharmacon | C-300576-05-0002 |
| miRIDIAN microRNA Human hsa-miR-511-5p - Mimic, 2 nmol | Dharmacon | C-300752-03-0002 |
| miRIDIAN microRNA Human hsa-miR-519c-3p - Mimic, 2 nmol | Dharmacon | C-300786-03-0002 |
| miRIDIAN microRNA Human hsa-miR-943 - Mimic, 2 nmol | Dharmacon | C-301275-01-0002 |
| miRIDIAN microRNA Human hsa-miR-200c-5p - Mimic, 2 nmol | Dharmacon | C-301070-01-0002 |
| miRIDIAN microRNA Human hsa-miR-944 - Mimic, 2 nmol | Dharmacon | C-301276-01-0002 |
| miRIDIAN microRNA Human hsa-miR-106a-3p - Mimic, 2 nmol | Dharmacon | C-301159-01-0002 |
| miRIDIAN microRNA Human hsa-miR-374b-5p - Mimic, 2 nmol | Dharmacon | C-301250-01-0002 |
| miRIDIAN microRNA Human hsa-miR-148b-5p - Mimic, 2 nmol | Dharmacon | C-301087-01-0002 |
| miRIDIAN microRNA Human hsa-miR-519d-3p - Mimic, 2 nmol | Dharmacon | C-300812-05-0002 |
| miRIDIAN microRNA Human hsa-miR-631 - Mimic, 2 nmol | Dharmacon | C-300957-01-0002 |
| miRIDIAN microRNA Human hsa-miR-592 - Mimic, 2 nmol | Dharmacon | C-300916-01-0002 |
| miRIDIAN microRNA Human hsa-miR-519b-3p - Mimic, 2 nmol | Dharmacon | C-300791-05-0002 |
| miRIDIAN microRNA Human hsa-miR-93-3p - Mimic, 2 nmol | Dharmacon | C-301207-01-0002 |
| miRIDIAN microRNA Human hsa-miR-340-5p - Mimic, 2 nmol | Dharmacon | C-301081-01-0002 |
| miRIDIAN microRNA Human hsa-miR-508-3p - Mimic, 2 nmol | Dharmacon | C-300848-05-0002 |
| miRIDIAN microRNA Human hsa-miR-374a-5p - Mimic, 2 nmol | Dharmacon | C-300681-05-0002 |
| miRIDIAN microRNA Human hsa-miR-520d-3p - Mimic, 2 nmol | Dharmacon | C-300816-05-0002 |
| miRIDIAN microRNA Human hsa-miR-520c-3p - Mimic, 2 nmol | Dharmacon | C-300803-05-0002 |
| miRIDIAN microRNA Human hsa-miR-520b - Mimic, 2 nmol | Dharmacon | C-300797-03-0002 |
| miRIDIAN microRNA Human hsa-miR-153-3p - Mimic, 2 nmol | Dharmacon | C-300615-07-0002 |
| miRIDIAN microRNA Human hsa-miR-519a-3p - Mimic, 2 nmol | Dharmacon | C-300836-05-0002 |
| miRIDIAN microRNA Human hsa-miR-1287-5p - Mimic, 2 nmol | Dharmacon | C-301341-00-0002 |
| miRIDIAN microRNA Human hsa-miR-373-3p - Mimic, 2 nmol | Dharmacon | C-300680-03-0002 |
| miRIDIAN microRNA Human hsa-miR-302a-3p - Mimic, 2 nmol | Dharmacon | C-300653-05-0002 |
| miRIDIAN microRNA Human hsa-miR-302b-3p - Mimic, 2 nmol | Dharmacon | C-300669-05-0002 |
| miRIDIAN microRNA Human hsa-miR-509-3p - Mimic, 2 nmol | Dharmacon | C-300849-07-0002 |
| miRIDIAN microRNA Human hsa-miR-210-3p - Mimic, 2 nmol | Dharmacon | C-300565-03-0002 |
| miRIDIAN microRNA Human hsa-miR-302c-3p - Mimic, 2 nmol | Dharmacon | C-300671-05-0002 |
| miRIDIAN microRNA Human hsa-let-7a-2-3p - Mimic, 2 nmol | Dharmacon | C-301297-00-0002 |
| miRIDIAN microRNA Human hsa-miR-224-5p - Mimic, 2 nmol | Dharmacon | C-300581-07-0002 |
| miRIDIAN microRNA Human hsa-miR-302d-3p - Mimic, 2 nmol | Dharmacon | C-300672-05-0002 |
| miRIDIAN microRNA Human hsa-miR-520e - Mimic, 2 nmol | Dharmacon | C-300772-03-0002 |
| miRIDIAN microRNA Human hsa-miR-520a-3p - Mimic, 2 nmol | Dharmacon | C-300788-03-0002 |
| miRIDIAN microRNA hsa-miR-20a-3p mimic, 2 nmol | Dharmacon | C-301022-01-0002 |
| miRIDIAN microRNA Human hsa-miR-498 - Mimic, 2 nmol | Dharmacon | C-300771-03-0002 |
| miRIDIAN microRNA Human hsa-miR-665 - Mimic, 2 nmol | Dharmacon | C-301246-01-0003 |
| miRIDIAN microRNA Mimic Negative Control #1, 5 nmol | Dharmacon | CN-001000-01-05 |

### Confirmation assay

HeLa cells were reverse transfected with 10 nM of the indicated miRNAs using Lipofectamine RNAiMAX according to manufacturer’s instructions. 72h after transfection the cells were trypsinized, harvested in 1X flow cytometry buffer (1X PBS, 1mM EDTA, 2% FBS) and analyzed for green and red fluorescence by flow cytometry using a BD FACSymphony instrument (BD-Biosciences).

### Western blots and antibodies

After incubation for 72h, transfected cells were washed once with 1X PBS, lysed directly in wells with 100-150µl of SDS Lysis buffer (62.5mM Tris-Cl, pH 6.8, 2% SDS) and sonicated (amplitude of 35%; 10 pulses with 2 seconds ON and 0.5 seconds OFF). The lysates were centrifuged for 2 minutes and protein concentrations quantitated by BCA Protein Assay (Pierce). Lysates (10 µg of protein/well) were run on 20-well 4-12% NuPage BisTris midi gels in 1X MES buffer (ThermoFisher Scientific) and transferred to polyvinylidene difluoride (PVDF) membranes using 25 mM Tris-HCl pH 7.6, 192 mM glycine, and 10% methanol. Membranes were blocked in 5% nonfat dried milk in Tris-buffered saline [pH 7.4], 0.05% Tween 20 (TBST) and then incubated with primary antibodies as follows: mouse monoclonal anti-E6AP (clone E4) at 1:1000 dilution (sc-166689 from Santa Cruz Biotechnology); rabbit monoclonal anti-cleaved PARP (Asp214) (D64E10 XP® #5625 from Cell Signaling Technologies); mouse monoclonal anti-HPV18-E6 (clone G-7) at a 1:250 dilution (sc-365089 from Santa Cruz Biotechnology); mouse monoclonal anti-p21 Waf1/Cip1/CDKN1A (clone 187) at a 1:1000 dilution (sc-817 from Santa Cruz Biotechnology); and mouse monoclonal anti-HPV18-E7 (clone F-7) at a 1:250 dilution (sc-365035 from Santa Cruz Biotechnology). Membranes were washed in TBST and incubated with horseradish peroxidase (HRP)-conjugated anti-mouse antibody at a 1:1,000 dilution (R1005 from Kindle Biosciences). The primary antibodies against β-Actin and P53 were directly conjugated to HRP and were used as follows: mouse monoclonal anti-β-Actin-HRP at a 1:50,000 dilution (A3854 from Millipore Sigma); and goat polyclonal anti-p53-HRP at a 1:3,000 dilution (HAF1355 from R&D systems). All blots were developed using KwikQuant Ultra Digital-ECL^TM^ Substrate Solution and processed using KwikQuant^TM^ Imager (Kindle Bioscience LLC). Western blots were quantified using the Image Studio Lite program (LI-COR Biotechnologies).

### RNA isolation and qRT-PCR

At 72h post-transfection, total RNA was isolated using the Nucleospin RNA kit (Takara Bio) according to the manufacturer’s instructions. Samples were quantified using NanoDrop™ Lite Spectrophotometer and equal amounts of RNA (500ng) were reverse transcribed using the high-capacity cDNA reverse transcription kit (Applied Biosystems). Quantitative real-time PCR (qRT-PCR) was performed in an Applied Biosystems ABI 7500 fast sequence detection system using TaqMan Fast Advanced Master Mix (ThermoFisher Scientific), and the following TaqMan custom gene expression assays for HPV18-E6E7: HPV18-E6E7 FWD 5’-CAACCGAGCACGACAGGAA-3’; HPV18-E6E7 PROBE 5’-AATATTAAGTATGCATGGACCTAAGGCAACATTGCAA-3’; and HPV18-E6E7 REV 5’-CTCGTCGGGCTGGTAAATGTT-3’ and for HPV16-E6E7: HPV16-E6-FWD 5’-AGGAGCGACCCGGAAAGT-3’; HPV16-E6-PROBE 5’-ACCACAGTTATGCACAGAGCTGCAAACAA-3’; and HPV16-E6-REV 5’-CACGTCGCAGTAACTGTTGCTT-3’. All reactions were normalized using a TaqMan gene expression assay for PPIA (Hs99999904_m1 ThermoFisher Scientific).

### Luciferase assays

C33A cells were transfected in 12-well plates with 10 nM of miRNAs as described above. After 24h, the miRNA transfection mix was removed and cells were transfected with 0.5 µg of either plasmid pGL4-LCRHPV16 (p6239, HPV16 nt 7000-85) (49) or pGL4-LCRHPV18 (p5194, HPV18 nt 6943-105) (50), containing the firefly luciferase ORF under the control of the HPV16 and HPV18 LCRs respectively, together with 0.5µg of a plasmid expressing the renilla luciferase under the control of the human β-actin promoter using Lipofectamine 3000 (ThermoFisher Scientific) according to manufacturer’s protocol. At 48h post-transfection of plasmids (72 hours after miRNA transfection), cells were lysed in the plate with 250 μl of 1x Passive Lysis Buffer and luciferase activity was determined using the Dual-Luciferase Reporter Assay System (Promega) according to the manufacturer’s instructions. Luciferase intensities were measured using a SpectraMax L luminescence microplate reader (Molecular Devices). Firefly luciferase readings were normalized to the respective renilla luciferase readings.

### Apoptosis assays

HeLa cells were reverse transfected with 10 nM of the different miRNAs either alone or combined with 20 nM of siRNAs directed to p53 (Dharmacon, D-003329-26: 5’-GCUUCGAGAUGUUCCGAGA-3’) or its C911 version (sense, 5’-GCUUCGAGUACUUCCGAGA-UU-3’) using Lipofectamine RNAiMAX according to manufacturer’s instructions. After incubation for 72h, attached cells were trypsinized and combined with all non-adherent cells in the supernatant and PBS washes. The cells were stained with Annexin V using the Dead Cell Apoptosis Kit with Annexin V Alexa Fluor™ 488 & Propidium Iodide (PI) (ThermoFisher Scientific #V13241 and #V13245). Stained cells were analyzed by flow cytometry using a BD FACSCanto flow cytometer (BD Biosciences).

### Figures and statistics

All graphics and statistical analyses were conducted using Prism 8 (GraphPad). When the values corresponding to mock transfections were used to normalize other values obtained in an experiment, the mock values were not included in the corresponding graphic since invariably their mean is equal to one and their standard deviation is zero. Figures were assembled using Affinity Photo (Affinity).

## Acknowledgements

We thank other members of the Howley laboratory for helpful discussions and suggestions. All plasmids generated in the Howley laboratory that are described in this manuscript are available through Addgene. GMN, JAS and PMH are co-inventors on a patent filed by Harvard on the therapeutic use of miRNAs in hrHPV positive cancers. This work has been supported by NIH grant R35CA197262 to P.M.H.

